# Subgenual anterior cingulate cortex controls sadness-induced modulations of cognitive and emotional network hubs

**DOI:** 10.1101/163709

**Authors:** Juan P. Ramírez-Mahaluf, Joan Perramon, Begonya Otal, Pablo Villoslada, Albert Compte

**Affiliations:** Institut d’Investigacions Biomèdiques August Pi i Sunyer (IDIBAPS), Barcelona, Spain

**Keywords:** depression, functional connectivity, graph, prefrontal cortex, frontopolar cortex

## Abstract

The regulation of cognitive and emotional processes is critical for proper executive functions and social behavior, but its specific mechanisms remain unknown. Here, we addressed this issue by studying with functional magnetic resonance imaging the changes in network topology that underlie competitive interactions between emotional and cognitive networks in healthy participants. Our behavioral paradigm contrasted periods with high emotional and cognitive demands by including a sadness provocation task followed by a spatial working memory task. We hypothesized that this paradigm would enhance the modularity of emotional and cognitive networks and reveal the hub areas that regulate the flow of information between them. By applying graph analysis methods on functional connectivity between 20 regions of interest in 22 participants we identified two main brain network modules, one cognitive and one emotional, and their hub areas: the left dorsolateral prefrontal cortex (dlPFC) and the left medial frontal pole (mFP). These hub areas did not modulate their mutual functional connectivity following sadness but they did so through an interposed area, the subgenual anterior cingulate cortex (sACC). Our results identify dlPFC and mFP as areas regulating interactions between emotional and cognitive networks, and suggest that their modulation by sadness experience is mediated by sACC.

## Introduction

Emotion and cognition are central to the quality and range of everyday human experience^1^. The question of how emotion and cognition interact is re-emerging motivated by advances in functional neuroimaging techniques and computational tools^2–6^.

Negative affect and anxiety are known to impair working memory (WM) performance^7–11^, which has been associated in neuroimaging studies with deactivation of cortical areas typically associated with WM in the prefrontal and parietal cortices^7, 12, 13^ and with the inverse activation of ventral regions typically associated with emotional processing^7, 8, 12, 13^. However, there is a strong integration between the “emotional brain” and the “cognitive brain” in most daily activities^3, 14, 15^ so that this segregation is generally blurry except for in the most extreme conditions such as in depressed patients^16^. Inducing recapitulation of a sadness experience in healthy subjects achieves a pattern of brain activations consistent with the strong opposition between emotional and cognitive circuits observed in depressed patients^17, 18^. Dorsal areas are typically deactivated (dorsolateral prefrontal cortex (dlPFC), dorsal anterior cingulate cortex (dACC) and parietal cortex) and ventral and subcortical areas are activated (insula, orbitofrontal cortex (OFC), subgenual anterior cingulate cortex (sACC), amygdala, hippocampus), while some other areas may play the role of interconnecting these two networks (basal ganglia, thalamus, rostral ACC)^17, 18^. At present, it is unclear whether these integration-segregation dynamics occur diffusely between all these participating areas, or whether there are specific areas that channel primarily these interactions. Deep brain stimulation (DBS) effectiveness in treatment-resistant depression but only at very specific points of the cingulo-frontal bundle suggests the presence of key nodes in this distributed network^19–21^. Here, we study the topology of sadness interactions within these brain circuits using network analysis of blood-oxygen level dependent (BOLD) time series with graph theory^2, 4, 22^, while participants engage in cognitive tasks in control and sadness conditions.

We hypothesized that a paradigm with a strong conflicting emotional and cognitive demand would enhance the modularity of emotional and cognitive brain networks in healthy participants and thus reveal the cortical areas that act as network hubs, which are critical for regulating the flow and integration of information between communities^23^. We collected functional magnetic resonance imaging (fMRI) data from 22 healthy subjects performing this task. Based on these activation patterns we extracted 20 regions of interest (ROI) on a subject-by-subject basis and we computed the correlations between fMRI time series in pairs of ROIs, obtaining the matrix of functional connectivity for each subject on which we applied network measures from graph theory^24^. We found that the dlPFC acted as connector hub of the cognitive subnetwork, and medial fronto-polar cortex (mFP) was the connector hub of the emotional subnetwork, but they both interacted via the sACC in the emotional subnetwork, and these connectivity patterns were associated with the intensity of the sadness experienced by the participants.

## Methods

### Participants

Twenty-two healthy participants (28.9 ± 3.9 years of age, mean ± s.d., 10 males) without any psychiatric, neurological or medical illness were recruited. All participants were screened with the Mini-International Neuropsychiatric Interview (M.I.N.I.) to specifically ensure the absence of any ICD-10 psychiatric disorders^25^ as well as those using psychoactive medications. All subjects were screened with Charlson comorbidity index^26^. All volunteers had normal or corrected-to-normal vision and were right-handed, native Spanish speakers. The study was carried out in accordance with ethical guidelines and deontological criteria established by the Hospital Clínic of Barcelona, as approved by its Ethics Committee of Clinical Research (Ref: 2009/4886) and written informed consent was obtained from all participants.

### Experimental Design

The study was composed of two different behavioral paradigms run immediately in sequence in the scanner (Fig. 1).

**Figure 1:**
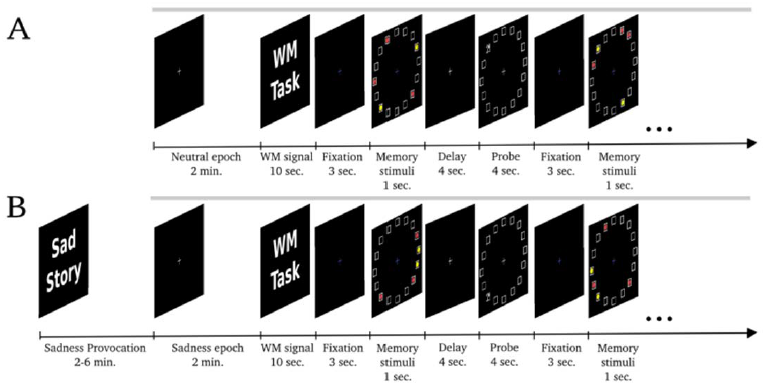
The two behavioral paradigms: Neutral-WM1 and Sadness-WM2. **A.**The first paradigm is composed of two tasks: a neutral epoch followed by 20 trials of spatial working memory with a filtering component. **B.**In the second paradigm, the participants underwent a sadness provocation task. When maximal sadness was achieved, participants closed their eyes for a 2-min scan, which was then followed by another 20 trials of spatial working memory with a filtering component. Note that in both paradigms the same stimuli were presented, the only difference being the sadness induced before in the second paradigm. Gray lines mark scanner acquisition periods (320 s).

In the first paradigm (Neutral-WM1), subjects were first instructed to rest in a “neutral emotional state” while keeping their eyes closed in a 2-min neutral-epoch scan. Following this resting condition, participants engaged in a spatial working memory task (WM1) with a filtering component^27^. In each of 20 working memory trials, 5 bright stimuli were presented on a black screen for 1 s, while participants were instructed to maintain fixation on a central cross. Stimuli appeared at random locations of equal eccentricity (6 deg. of visual angle) on a grid of 16 possible positions^27^. Of these stimuli, three were red dots and two were yellow dots. Participants were instructed to remember only the position of the red dots. Stimuli were followed by a dark screen during a delay of 4 s and then a probe stimulus was displayed for 4 s at one grid location (Fig. 1). At that point, participants pressed one of two buttons with the index or middle finger of their right hand to indicate whether one of the red dots had been presented at the location indicated by the probe or not, respectively.In 25% of the trials, the probe appeared at a location previously occupied by a yellow dot. In these trials, an error response of the participants (yes response) indicated a failure to apply the filtering component of the task. We define this type of error as a cognitive inhibition error^27, 28^.

In the second paradigm (Sadness-WM2), participants performed an emotional task, the sadness provocation task (SP)^29^. All subjects had prepared in advance a short autobiographical narrative of personal events in which they felt particularly sad, e.g. sad experiences most commonly centered on the loss of relatives, friends, or significant relationships. Before scanning, the narrative texts were presented on the screen and subjects were asked to generate a state of sadness comparable to that originally experienced. After the maximum mood intensity was achieved, participants pressed a button, closed their eyes and were instructed to stop visualizing, thinking, or ruminating on the text and to focus on their feelings of sadness while we acquired fMRI data in a 2-min sadness-epoch scan. Following the sadness epoch, and without stopping the scanner, participants performed a second spatial working memory task (WM2) with filtering component, following the description in the first paradigm. After the scan session, participants reported their subjective rating of the sadness experienced on a 0–7 point scale (5.6 ± 0.17, mean ± SEM)^30^.

All participants trained for all tasks before the scanner session. We did not reverse the order of the paradigms to avoid that sadness affected the Neutral-WM1 paradigm.

### Behavioral analysis

Participants were divided into two groups depending on their sadness intensity: “*high-sadness* group” were those whose subjective sadness rating ranked above the overall mean (*n*=12, mean=6.14, median=6, range 6-7) and “*low-sadness* group” were those participants whose rating ranked below the mean (*n*=10, mean=4.85, median=5, range 4-5.5). Two participants were excluded from the behavioral analysis because they reported difficulty in distinguishing the color of the dots during the WM tasks (1 from the *high-sadness* group, and 1 from the *low-sadness* group). We measured WM performance with the fraction of errors, the reaction times and the fraction of cognitive inhibition errors for each subject in WM1 and WM2.

### fMRI acquisition

Brain images were acquired on a 3 Tesla TimTrio scanner (Siemens, Erlangen, Germany) using the 8-channel phased-array head coil supplied by the vendor. A custom-built head holder was used to prevent head movement, and earplugs were used to attenuate scanner noise. High-resolution three-dimensional T1-weighted magnetization prepared rapid acquisition gradient echo (MPRAGE) images were acquired for anatomic reference (TR=2200 ms, TE=3 ms, FA=7º, 1.0 mm isotropic voxels). A T2-weighted scan was used in order to detect possible pathological features (TR=3780 ms, TE=96 ms, FA=120º, voxel size 0.8x0.6×3.0mm, 3.0mm thick, 0.3mm gap between slices, 40 axial slices). Functional data were acquired using a gradient-echo echo-planar pulse sequence sensitive to blood oxygenation level-dependent (BOLD) contrast (TR=2000 ms, TE=30 ms, FA=85º, 3.0mm isotropic voxels, 3.0 mm thick, no gap between slices). Presentation® software and data acquisition were synchronized to stimulus pulse sent by the scanner. Participants were requested to avoid moving during the whole MRI scan. The total duration of uninterrupted scanning time during each of the two behavioral paradigms (Fig. 1) was 320 s, i.e. 160 volumes.

### fMRI data analysis

Preprocessing and statistical analysis were carried out with SPM8 (Wellcome Trust Centre for Neuroimaging, http://www.fil.ion.ucl.ac.uk/spm). The images were manually aligned along the anterior commissure-posterior commissure line. Preprocessing included the realignment of the scans for motor correction and the normalization to the Montreal Neurological Institute (MNI) template (interpolating to 3 mm cubic voxels). For GLM analyses we further applied spatial smoothing with a Gaussian kernel of 10 mm (we also tried Gaussian kernels of 8mm and 6mm, which did not affect the GLM results). For the functional connectivity analysis the spatial smoothing was not applied because we averaged all voxels within each ROI prior to the connectivity analyses.

A random-effect, epoch-related statistical analysis was performed in a two-level procedure. At the first level, a general linear model (GLM) was estimated by using regressors for each instruction condition (before WM trials), neutral and sadness epochs, and fixation period, memory stimulus, delay period and probe stimulus (Fig. 1). Regressors were convolved with the canonical hemodynamic response function in SPM8. The data were high-pass filtered (128 s cutoff) to remove low-frequency drifts. Images from contrasts of interest for each participant were used in a second-level analysis, treating participants as a random effect.

A paired sample t-test was used to investigate the resulting statistical maps for the contrast *delay-fixation* in WM1 and WM2. The voxel significance was evaluated in a whole-brain analysis testing the global null hypothesis that *delay-fixation* showed no significant activation. This analysis was corrected for multiple comparisons (false discovery rate (FDR), *P* < 0.05) and it identified 10 different cortical areas implicated in cognitive processing in this task (Table 1).

**Table 1:**
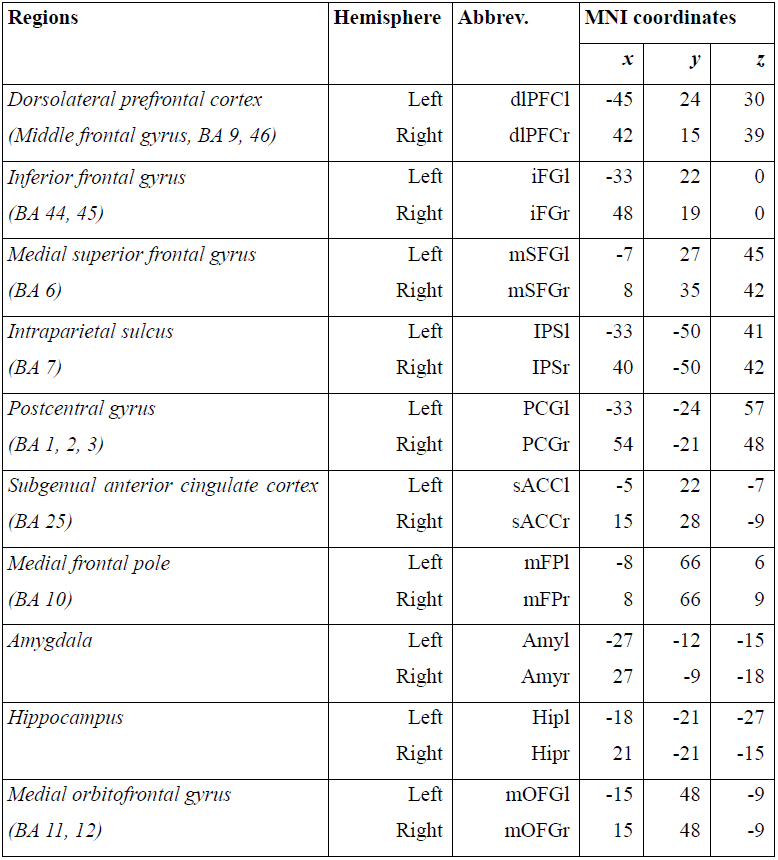
Regions identified in the task-based analyses. MNI coordinate system. BA = Broadmann area

Subsequently, a paired sample t-test was used to statistically assess the difference between delay activity in WM1 and WM2. A mask was created including the activated areas in both WM1 and WM2 in order to compare the difference in the level of activation between WM1 and WM2. The voxel significance was evaluated in the mask testing the global null hypothesis that *delay*_*WM1*_-*delay*_*WM2*_ did not show significant activation. This analysis was corrected for multiple comparisons in the working memory mask (false discovery rate (FDR), *P* < 0.05).

We used a 2-factor ANOVA to statistically assess the interaction between delay activity in the two tasks (*delay*_*WM1*_ and *delay*_*WM2*_) and sadness intensity (*high-sadness* and *low-sadness* groups). The voxel significance was evaluated in a mask testing the global null hypothesis that *delay*_*WM1*_ -*delay*_*WM2*_ and *high-sadness* -*low-sadness* groups did not show significant interactions.

We further used a 2-factor ANOVA to statistically assess the interaction between brain activity in the passive conditions (*Neutral* and *Sadness*) and sadness intensity (*high-sadness* and *low-sadness* groups). Voxel significance was evaluated in a whole-brain analysis testing the global null hypothesis that the interaction between *Sadness/Neutral* and *high-sadness/low-sadness* groups was not significant. This analysis led us to identify 9 different cortical and subcortical areas implicated in emotional processing in this task (*P*_*unc*_ < 0.05) (Table 1).

For the sACC, which shows significant interaction (small volume correction, 5 mm square at −6 21 −9, FWE-corrected, *p* = 0.038), a paired sample t-test was used to statistically assess the difference between *Sadness* and *Neutral* epochs in the *high-sadness* group.

### Functional connectivity

Cognitive and emotional regions of interest (ROIs) were determined from global analysis as indicated in Results (see list of ROIs in Table 1).

Some areas known to be related to emotional processing did not survive the correction by multiple comparisons; medial orbitofrontal gyrus left, subgenual anterior cingulate cortex right, medial frontal pole bilateral, hippocampus bilateral and amygdala bilateral. Medial orbitofrontal gyrus right did not show significant activation and was added as the laterally symmetric counterpart of the medial orbitofrontal gyrus left. The locations of the emotional ROIs taken around the peak activations of our contrasts were consistent with coordinates in the literature^29, 31–35^. Each ROI was defined as a 5 × 5 × 5 voxel cube centered around the detected peak activations (for coordinates see Table 1).

The ROI signals were obtained by linear detrending preprocessed data without spatial smoothing for each voxel, and then by averaging across all voxels within the ROI. We removed covariations common to all ROIs by applying a signal regression^36, 37^. This procedure removes global fluctuations related to physiological artifacts such as heart rate, respiration, and scanner noise that are seen throughout the brain artificially, but it could also introduce spurious anticorrelations (See Discussion)^36, 38, 39^.

To focus on temporal fluctuations of the BOLD signal not related to the imposed structure of the paradigms (two tasks (SP/resting and WM), change of task in a time scale of 2 min) ROI signals were band-pass filtered in the range 0.018-0.26 Hz (i.e. maintaining temporal fluctuations in time scales from 4 seconds to 1 minute). Functional connectivity between areas was computed with the Pearson correlation coefficient between the 320-sec signals for each pair of ROIs in our database (Table 1), separately for each of the two behavioral paradigms (Fig. 1).

We tested the functional significance within our task of correlations between ROIs identified as hubs of the network (see below) with ANOVA tests. For sACCl-dlPFCl and sACCl-mFPl correlations, we ran a 3-factor ANOVA tests with the factors: paradigm (*Neutral-WM1* vs. *Sadness-WM2)*, sadness intensity (*high-sadness* vs. *low-sadness* groups) and participant identity as a random factor. The interaction between sadness intensity and paradigm was significant for both connections (false discovery rate (FDR), *P* < 0.05; *p=*0.0006 for sACCl-dlPFCl; *p* = 0.0434 for sACCl-mFPl), so we separated the data for each group and we performed a paired sample t-test comparing correlations for different sadness groups and paradigms.

### Graph analysis

For each subject and behavioral paradigm, the correlation matrix between our ROIs is the adjacency matrix of the weighted graph that represents the corresponding brain network^2^. The symmetrical adjacency matrix resulting from our undirected graph was characterized for having positive and negative weights. We used algorithms adapted to this type of data using standard graph theory methods on Matlab (Brain Connectivity Toolbox developed by O. Sporns, Indiana University, Bloomington, IN; (https://sites.google.com/site/bctnet/)^24, 40^. We calculated the community structure from the mean correlation matrix across subjects. To identify the best partition in modules (communities), we quantified its modularity by a quality function *q*, which we optimized. *q* is a scalar value between –1 and 1 that measures the density of links inside communities as compared to links between communities. Large values of *q* reflect more segregation, or equivalently, decreased integration, between different communities. The best partition is defined as the set of communities with the largest modularity *q*. The modularity analysis has two free parameters (resolution parameters γ***+*** and γ–) that allow weighing differently the positive and negative correlations in the connectivity, and this has an impact in the community structure that the method identifies. We used resolution parameters (γ***+*** = 1, γ– = 1), and (γ***+*** = 1, γ– = 0.75) in our analyses (Results)^41, 42^.

Once we calculated the community structure, we measured the degree and the participation coefficient of each node (i.e. ROI in Table 1), and the global efficiency of each network community. The degree of a node is the number of connections to that node. The degree has a straightforward neurobiological interpretation: nodes with a high degree are interacting, structurally or functionally, with many other nodes in the network. In our weighted graph, we validated a connection if its corresponding absolute weight exceeded a pre-defined threshold (range 30%-45%) relative to the absolute strongest correlations. In the results, we consider high-degree areas when their degree is greater than the network mean degree plus one standard deviation. These areas are candidates to be defined as hubs of the networks, as previously argued in the literature^23^. The Participation coefficient is a measure of diversity of intermodular connections of individual nodes, it compares the degree of a given node to the number of connections within its own subnetwork. This measure requires a previously determined community structure (see above)^23^. Global efficiency (GE) measures the average strength of the shortest paths in the network and can be interpreted as the overall “efficiency of communication” minimizing the cost of communication over the most direct paths in the networks. Global efficiency requires as inputs a measure of node dissimilarity, or the “cost” of a connection; which we defined as the inverse of the functional connection weight. We calculated GE for each community, separately. Since within each community most of the functional connectivity weights are positive, negative weights were set to zero for this analysis^24^.

Statistical significance of group differences was assessed with permutation tests, where we permuted randomly the assignment of data to each group and we repeated the measure 1,000 times. We reported a significant difference if the difference between the actual groups was larger that 95% of the samples generated randomly (*p* < 0.05).

## Results

We recorded fMRI brain activity in 22 participants while they engaged in two identical working memory tasks (WM1, WM2), separated by a period in which a sadness state was induced by remembering a previously identified biographical sketch (Fig. 1). After the scan session, participants reported their subjective rating of the sadness intensity. We sought to identify the functional changes induced by the sadness state in brain networks engaged in regulating the interaction between cognition and emotion.

### Behavioral analysis

Across participants, the mean number of error trials, of cognitive inhibition error trials (see *Experimental design*), and the mean reaction times did not change significantly from the working memory session before sadness induction (WM1) to the working memory session after sadness induction (WM2) (Table 2, paired sample t-test: *p* = 0.5, *p* = 0.6, *p* = 0.85, respectively, n=20).

**Table 2:**
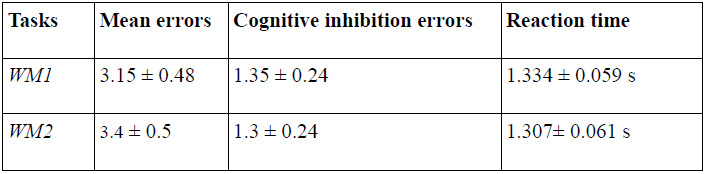
Behavioral measures. Indicated are parameters of behavioral responses in the 20-trial WM tasks of the Neutral-WM (WM1) and Sadness-WM (WM2) paradigms. We report mean ± SEM, n=20.

As defined in our analysis, we split participants according to their subjective report on sadness intensity (see *Behavioral analysis*). We found that participants in the *high-sadness* group diminished their WM performance following sadness provocation (Fig. 2A, Table 3, 3-way ANOVA with factors high-sadness/low-sadness, WM1/WM2 and subject identity, n=20, *p* = 0.04 for the interaction between *high-sadness/low-sadness* groups and WM1/WM2, paired sample t-test *p* = 0.033 for WM1-WM2 errors in the *high-sadness* group, *p* = 0.37 for WM1-WM2 errors in the *low-sadness* group). This underscores the validity of this subjective report, so that we used it in the following to identify effects associated specifically with the experience of sadness.

**Table 3:** Mean number of errors in *high-sadness* and *low-sadness* groups. Mean number of errors in the 20-trial WM tasks of the Neutral-WM (WM1) and Sadness-WM (WM2) paradigms. We report mean ± SEM, n=11 (high-sadness), n=9 (low-sadness).

|  | ***High-sadness:***<br>**Mean errors** | ***Low-sadness:***<br>**Mean errors** |
| --- | --- | --- |
| *WM1* | $3.27 \pm 0.74$ | $3 \pm 0.62$ |
| *WM2* | $4.18 \pm 0.75$ | $2.44 \pm 0.5$ |

**Figure 2:**
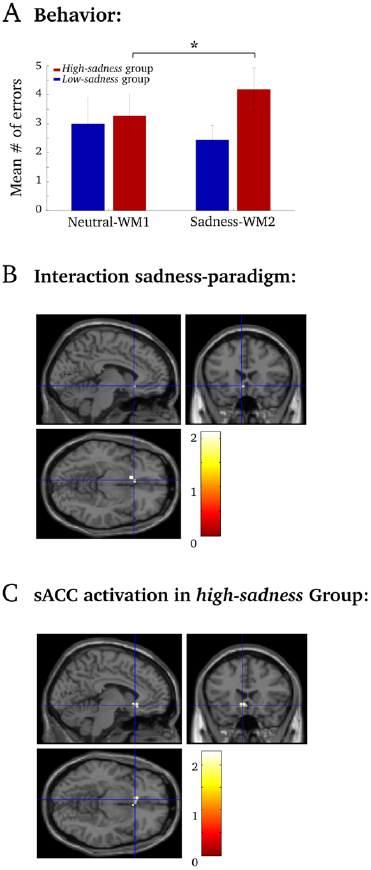
Sadness intensity disrupted WM performance and activated the sACCl. **A.**Mean number of error trials during working memory tasks: The *high-sadness* group presented more errors in WM2 relative to WM1. **B.**A significant interaction between *Sadness/Neutral* and *high-sadness/low-sadness* groups in sACCl (ANOVA T-contrast, 6-voxel cluster, peak activation at –6 21 –9. **C.**Significant sACCl activation during sadness relative to neutral states (Fig. 1) in the *high-sadness* group (20-voxel cluster, peak activation at –3 21 –9. The intersection of lines marks the peak cluster activation of **B** (–6 21 –9 coordinates).

### fMRI BOLD response during working memory

We first identified the cortical areas that are supporting the memory component of the working memory task in the cognitive network during WM1 and WM2. We conducted a whole brain analysis to find regions activated in a *delay*-*fixation* contrast (see *fMRI data analysis*). We found significant activation (False Discovery Rate, FDR, *p* < 0.05, Supplementary Fig. S1) in the cognitive areas: dorsolateral prefrontal cortex (dlPFC), intraparietal sulcus (IPS), medial superior frontal gyrus (mSFG), postcentral gyrus (PCG) and inferior frontal gyrus (iFG) (Supplementary Fig. S1A for WM1 and Supplementary Fig. S1B for WM2, for coordinates see Table 1). We tested changes in delay activity in these areas by applying a mask on a *delay*_*WM1*_-*delay*_*WM2*_ contrast (Supplementary Fig. S1C), and we found a significant reduction in activity during *delay*_*WM2*_ in all these areas (FDR, *p* < 0.05). However, we could not attribute this decrease unambiguously to sadness experience, as we could not find an interaction between the factors *high-sadness/low-sadness* and WM1/WM2 in none of these areas (2-way ANOVA, FDR *p* < 0.05, mask with areas in Supplementary Fig. S1A-B).

### fMRI BOLD activity related to sadness experience

We then looked for the anatomical regions activated during the *Sadness* epoch (Fig. 1). Previous studies have pointed to the sACC as an area involved in sadness processing.Across all participants, we did not find a significant activation in the sACC, or in any other area, in the *Sadness* epoch relative to the *Neutral* epoch (2-way ANOVA whole brain analysis with factors epoch and sadness intensity, T-contrast, FDR *p* < 0.05). Nevertheless, it is known that factors associated with individual differences at both neuroanatomical and behavioral levels may account for the difficulty in detecting sACC activation (Smith et al., 2011). We thus resorted to a region of interest (ROI) analysis, where we defined the sACC ROI (125-voxel cube, center in Table 1) based on available evidence from previous neuroimaging studies^29, 32, 33^. Using this ROI as a mask in the above analysis, we found a significant interaction between *Sadness/Neutral* epoch and *high-sadness/low-sadness* groups in the left hemisphere (sACCl, Fig. 2B, cluster of 6 voxels, peak activation at –6 21–9, 5-mm-square small volume SVC and FWE corrections, *p* = 0.038). Sadness provocation evoked an increase in the BOLD signal in the sACCl during *Sadness* compared with *Neutral* in the *high-sadness* group (Fig. 2C, cluster of 20 voxels, peak activation at –3 21 –9, SVC and FWE, *p* = 0.014). In other words, subjects who achieved an intense sadness state activated the left sACC.

Thus, the association of sACC activations with sadness reports was in contrast with its lack of statistical significance in FDR-corrected whole-brain *Neutral-Sadness* contrasts. Based on this result, we decided to define the network of areas putatively involved in sadness processing by lowering our statistical threshold (whole brain uncorrected tests at *p*_*unc*_ < 0.05) in a 2-factor ANOVA testing the interaction between the factors epoch (*Neutral* / *Sadness*) and sadness intensity (*high-sadness* / *low-sadness*). This analysis led us to identify 9 different bilateral cortical and subcortical areas implicated in emotional processing in this task (see Table 1).

### Community structure distinguishes emotional and cognitive networks

Based on these BOLD activations, consistent with previous literature^29, 31–35^ we thus defined a set of ROIs (Table 1) that would be presumably implicated in the regulation of cognitive and emotional task demands, and we set to determine how sadness experience was associated with functional changes in network topology.

For each pair of ROIs we estimated their functional connectivity as the linear dependence of the temporal fluctuations in the corresponding signals, as measured by the Pearsoncorrelation coefficient. This led us to define a symmetrical connectivity matrix containing the correlation coefficients between all possible pairs of ROIs. This matrix consists of positive and negative correlations (see *Functional connectivity* and *Graph analysis*). We obtained one such connectivity matrix independently for each subject, and we then averaged together these matrices to obtain a matrix of the averaged connectivities across participants. We applied graph-theoretic analyses by considering ROIs as nodes and the functional connectivity between each pair of ROIs as the corresponding edge.

We first asked if the pattern of connectivities defined subnetworks of areas that had distinct connectivity within and across subnetworks. This can be determined through a community detection algorithm that finds the assignment of nodes (ROIs) in communities (subnetworks) by maximizing the *modularity q* of the partition (see *Graph analysis*). This community detection algorithm applied to our experimental grand-average connectivity matrix identified two main communities that coincide with the results of our BOLD contrast analyses above: the cognitive module (areas mSFG, PCG, IPS, dlPFC and iFG, Supplementary Fig. S1), and the emotional module (areas sACC, medial Frontal Pole (mFP), medial orbitofrontal gyrus (mOFG), Amygdala (Amy) and Hippocampus (Hip)) that was related to sadness (Fig. 3A-B, Supplementary Fig. S2A-B). The pattern of correlations shows that these subnetworks interact with each other mainly through positive correlations (Fig. 3A-B, Supplementary Fig. S2A-B, red lines) and between them mainly through negative correlations (Fig. 3A-B, Supplementary Fig. S2A-B, blue dashed lines), as seen in the distributions of correlations (Figs. 3C and Supplementary Fig. S2C). The *modularity* was higher for *Sadness-WM2* than for *Neutral-WM1* (*q* = 0.431 vs. *q* = 0.412, permutation test, *p* = 0.008, Table 4), suggesting that the emotional and cognitive communities get more segregated following an episode of intense sadness. We confirmed this by applying the community detection algorithm to the average correlation matrices obtained separately for the *high-sadness* and the *low-sadness* groups in the paradigm *Sadness-WM2.* The community assignment of the different ROIs did not change based on sadness intensity (Fig. 3D-E), but the modularity *q* was indeed higher for the *high-sadness* group than for the *low-sadness* group (*q* = 0.443 vs. *q* = 0.405, permutation test, *p* < 0.0001, Table 4), confirming our hypothesis.

**Table 4:** Network-and community-level measures. Arrow denotes the direction of change, and items in bold are statistically significant. NS = non-significant. Significance (*p* value) calculated using permutation tests.

|  | *Sadness-WM2* /<br>*Neutral-WM1* | *high-sadness* /<br>*low-sadness* |
| --- | --- | --- |
| Modularity $Q$ ($g+ = g- = 1$) | ↑ (**$p = 0.008$**) | ↑ (**$p < 0.0001$**) |
| Modularity $Q$ ($g+ = 1, g- = 0.75$) | ↑ (**$p < 0.0005$**) | NS ($p = 0.14$) |
| Global efficiency (emotional) | ↑ (**$p = 0.016$**) | NS ($p = 0.72$) |
| Global efficiency (cognitive) | NS ($p = 0.27$) | NS ($p = 0.56$) |

**Figure 3:**
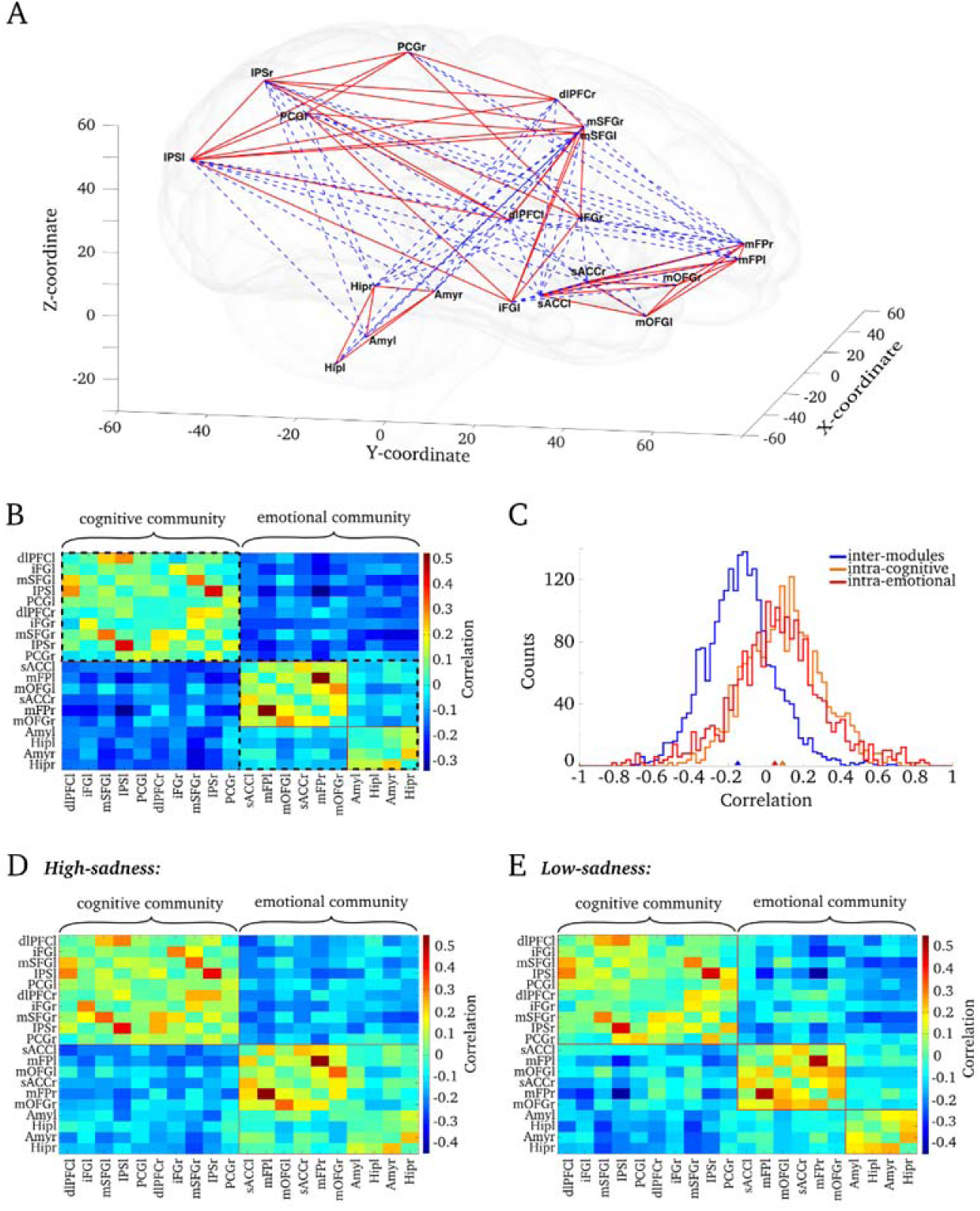
Cognitive and emotional communities for Sadness-Working memory. **A.**3D-graphical representation of the networks, the ROIs are located according to real-world coordinates. Mean significant correlations are plotted; positive correlations in red lines and negative correlations in blue dashed lines, shading brain for schematic purposes. **B.**Matrix of the mean correlations across subjects. The analysis identified two main modules, the cognitive and emotional communities separated by the dashed black line. Into the emotional community, two sub-communities were found (separated by the red line), corresponding to the emotional areas in the cortex and the limbic system (subcortical areas). **C.**Correlation distributions for all subjects. The correlations between cognitive and emotional modules (inter-modules), plotted in blue were mainly negative (mean ± SEM = –0.15 ± 0.004). The correlations within the cognitive module (intra-cognitive, plotted in orange, mean ± SEM =0.089 ± 0.005) and the correlations within the emotional module (intra-emotional, plotted in red, mean ± SEM = 0.048 ±0.009) were both positively biased. **D.**Matrix of the mean correlation for *high-sadness* subjects. The analysis identified two main modules, cognitive and emotional communities separated by the red line. **E.**Matrix of the mean correlation for *low-sadness* subjects. The analysis identified three modules, one cognitive and two emotional communities separated by the red line. Note that for the low-sadness group, but not for the high-sadness group, the algorithm was able to detect within the emotional community the two sub-communities reported for all subjects (cortical and subcortical areas).

When we had the community detection algorithm apply more weight to the positive correlations (see *Graph analysis*), it identified two sub-communities within the emotional community (Figs. 3B and Supplementary Fig. S2B, red line). These sub-communities corresponded to emotional areas in the cortex (sACC, mOFG and mFP) and the limbic system (Amy and Hip), respectively. Consistent with this substructure in the emotional community, the mean correlation within the emotional community was lower than within the cognitive community (Figs. 3C and Supplementary Fig. S2C; 2-way ANOVA, *p* = 0.044 and *p* = 0.0062, respectively). In these conditions, the *modularity q* in *Sadness-WM2* was higher than in *Neutral-WM1* (*q* = 0.352 vs. *q* = 0.332, permutation test, *p* < 0.0005, Table 4), but it was not significantly different between the *high-sadness* and *low-sadness* groups during *Sadness*-WM2 (*q* = 0.355 vs. *q* = 0.345, Table 4). This reflected the fact that the two emotional sub-communities enhanced their integration in the *high-sadness* group: the community detection algorithm was unable to distinguish the emotional sub-communities in the *high-sadness* group (Fig. 3D). Taken together, this suggests that sadness experience causes the segregation of cognitive and emotional networks, while at the same time promoting more integration between cortical emotional areas (sACC, OFC and mFP) and the limbic system (Amy and Hip).

We then turned to studying whether subnetworks changed their internal connectivity in the task. Global efficiency (GE) computes an estimate of the average inter-node distance within a given community, and it provides a measure of its integration within the network. A community with higher GE will have “shorter paths” between the nodes (with path distancedefined as the inverse of the functional connectivity between two nodes, see *Graph analysis*). We calculated GE separately for each cognitive and emotional network, and separately for *Sadness-WM2* and *Neutral-WM1.* We found significantly higher GE for the emotional network during the *Sadness-WM2* compared to the *Neutral-WM1* paradigms (*GE* = 0.091 and 0.082, respectively, permutation test, *p* **=** 0.016), but not for the cognitive network (Table 4). When we compared the *high-sadness* and *low-sadness* groups, we did not find a significant difference in the GE of either the emotional or the cognitive networks (Table 4).

These results suggest that intense sadness mostly affects the interaction between emotional and cognitive subnetworks (increases modularity q), increases the internal integration of the emotional module (increases GE), and has little effect on interactions within the cognitive subnetwork (stable GE).

### Hub identification and their modulation by strong emotional demands

The network-level properties studied above suggest a modulation in the interaction between the networks according to the emotional or cognitive demands. We wondered if specific areas (*hubs*) mediated these interactions. We investigated this by measuring, in each participant’s connectivity matrix, two network parameters for each node: the degree, and the participation coefficient (see *Graph analysis*). The degree of a node in the network is the number of connections it has to other nodes, and the participation coefficient compares this number of connections to the number of connections within the node’s own subnetwork. Nodes with a high degree and a high participation coefficient are known as connector hubs, and they are candidates to mediate interaction between subnetworks^23^. For these measures we considered strong absolute correlations, above a threshold of the absolute maximal correlation, for each subject. We identified connector hub nodes as those ROIs with degree one standard deviation above the network’s mean degree, and with participation coefficient above the network’s mean participation coefficient, following the criteria of previous studies^23^.

During the *Neutral-WM1* paradigm we identified 5 connector hub ROIs: IPSr, IPSl, dlPFCl, mFPl and mSFGl (Fig. 4A). On the other hand, during the *Sadness-WM2* paradigm we identified 3 connector hubs: IPSr, IPSl and mFPr (Fig. 4B). These connector hubs were consistently identified independently of the threshold applied to the correlation matrix (Supplementary Figs. S3A, S3B)). We noticed that the IPS and mFP were present in both task paradigms, while dlPFCl and mSFGl appeared only in the *Neutral-WM1* paradigm. The drop of dlPFCl from the list of connector hubs was significant, in that its degree decreased significantly from *Neutral-WM1* to *Sadness-WM2* (3-way ANOVA with factors: task paradigm, sadness intensity, and subject, threshold 35%, main effect of paradigm *p* = 0.014, all other effects and interactions not significant, Fig. 4C), independently of the applied threshold (range 30%-45%, Fig. 5A). The dlPFCl participation coefficient also showed a marginally-significant decrease (Fig. 4C, main effect of paradigm *p* = 0.08). However, the degree of the dlPFCl was not significantly modulated by the intensity of sadness (3-way ANOVA, threshold 35%, interaction between task paradigm and sadness intensity *p* = 0.55). Instead, we found that the degree of the mFPl was modulated by sadness intensity in a range of thresholds: it increased for the *high-sadness* group and it decreased for the *low-sadness* group relative to *Neutral-WM1* (3-way ANOVA, interaction between task paradigm and sadness intensity, *p* < 0.05, Fig 5B). These results identified dlPFCl as a connector hub in the cognitive subnetwork that reduced its coupling following sadness induction, and mFPl as a connector hub in the emotional subnetwork that increased its coupling specifically in those participants that experienced a stronger emotional state after sadness induction.

**Figure 4:**
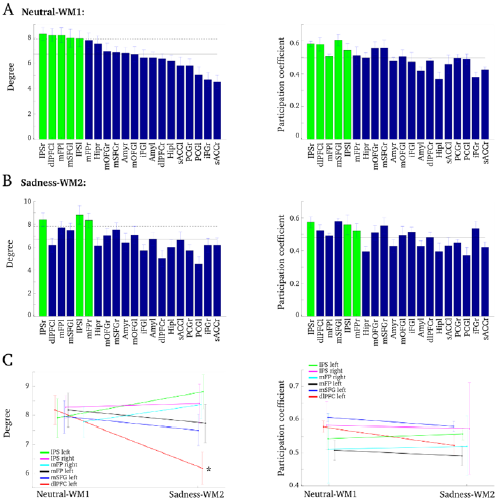
Hubs identification, dlPFCl decreases their degree after sadness. Degree *(left panels)* and participation coefficient *(right panels)* analysis applying a threshold of 35% to the correlation matrices of each subject. **A.**During the Neutral-WM1 paradigm 5 regions were identified as hubs (green bars): IPSr, IPSl, dlPFCl, mFPl and mSFGl. Their participation coefficients were above the mean, so they were classified as connector hubs. **B.**During the Sadness-WM2 paradigm 3 regions were identified as hubs (green bars): IPSr, IPSl, mFPr, which were classified as connector hubs. Note that the ordering of areas is the same as in **A**. **C.**The dlPFCl was the only hub that presented a significant decrease in the degree from Neutral-WM1 to Sadness-WM2and a marginally-significant decrease in participation coefficient. Error bars mark standard error of the mean.

**Figure 5:**
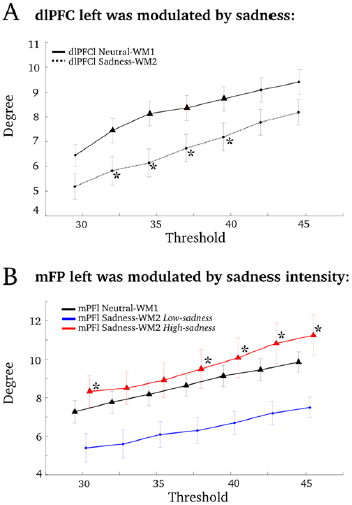
The dlPFCl was modulated across paradigms and mFPl was modulated by sadness intensity, robustly across thresholds of the correlation matrix. **A.**The decrease in the degree for the dlPFCl during Sadness-WM2 was stable across thresholds. Black asterisks mark significant main effect of paradigm (3-way ANOVA, *p* <0.05). **B.**The mFPl was the only hub that was modulated by the sadness intensity. It increased its degree in the *high-sadness* group and it decreased it in the *low-sadness* group. Black asterisks mark significant interaction between task paradigm and sadness intensity (3-way ANOVA, *p* <0.05).

We mark with a triangle when the corresponding region is classified as a hub among the rest of areas. Error bars mark standard error of the mean.

### Changes in functional connectivity underlie behavioral differences and hub modulations

We investigated the mechanism underlying the hub modulations described above by first testing if a change in the functional connectivity between the 2 connector hub nodes, dlPFCl and mFPl, could explain the modulations of their degree. We analyzed the change in the correlation between these 2 hub nodes for each participant, task paradigm and sadness-intensity groups. The functional connectivity between dlPFCl and mFPl did not present either a main effect of task epoch (*Neutral-WM1 vs. Sadness-WM2*, 3-way ANOVA, *p* = 0.48) or an interaction between *high-sadness/low-sadness* and *Neutral-WM1/Sadness-WM2*, (3-way ANOVA, *p* = 0.12). Direct interactions between the 2 hub nodes were thus not a mechanism supported in our data for the network modulation operated by the sadness state.

Then, we looked for other nodes that could mediate the modulation of the connector hubs. We analyzed the change in correlations (as a measure of functional connectivity) between the 2 connector hub nodes (mFPl and dlPFCl) and all other network areas for each participant, task paradigm and sadness-intensity groups. We thus tested a total of 19 pairwise correlations for each connector hub node, and we corrected our tests for the multiple comparison problem by controlling the false discovery rate (FDR) at a levelα=0.05.

The functional connectivity between dlPFCl and sACCl presented a significant interaction between *high-sadness/low-sadness* and *Neutral-WM1/Sadness-WM2*, (3-way ANOVA, *p* = 0.0006, *p* _(FDR-corr)_ = 0.036). The correlations between dlPFCl and sACCl became more negative after sadness provocation only in the *high-sadness* group (Fig. 6A, paired sample *t*-test *p*=0.0001 for *high-sadness* and *p*=0.49 for *low-sadness*). In other words, only the group reporting more intense sadness presented a stronger anticorrelation between sACCl and dlPFCl after sadness induction. This suggests that the interactions of dlPFCl with sACCl could be associated with the reduction in dlPFCl network degree following sadness induction (Figs. 4C and 5A).

**Figure 6:**
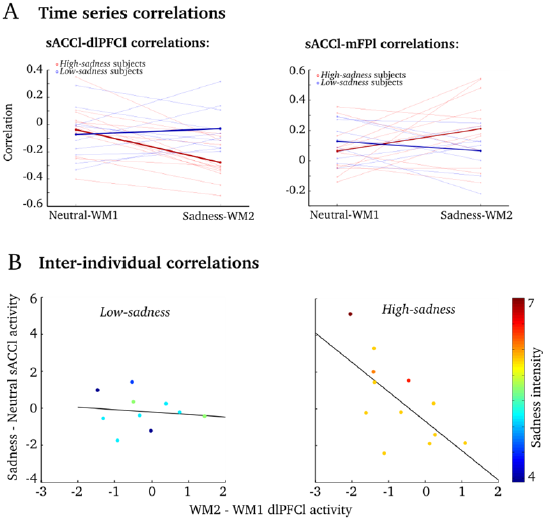
Sadness intensity increased the sACCl-dlPFCl anticorrelation and the sACCl-mFPl correlation. **A.**Subject-by-subject correlations for sACCl-dlPFCl and sACCl-mFP during Neutral-WM1 and Sadness-WM2. *High-sadness* subjects are plotted with red lines, *low-sadness* subjects with blue lines and the population averages are plotted in thick lines, respectively. *Left, High-sadness* subjects and not *low-sadness* subjects presented significant increased anticorrelations between sACCl and dlPFCl during Sadness-WM2 compared to Neutral-WM1. *Right,* sACCl and mFPl correlations presented a significant interaction between sadness intensity (*High-sadness* vs. *low-sadness* subjects) and paradigm (Neutral-WM1 vs. Sadness-WM2). **B.**For the high-sadness group (right) but not for the low-sadness group (left), we found a significant anticorrelation between sACCl (Sadness-Neutral) and dlPFCl (WM2-WM1) BOLD activity: the stronger the BOLD activity in sACCl during sadness, the weaker the BOLD activity in dlPFCl during WM2.

The anticorrelation between sACCl and dlPFCl was in addition related to the functional activation of sACCl (Fig. 2B-C) and deactivation of cognitive areas (Supplementary Fig. S1C) as we found a significant inter-individual anticorrelation between the contrast *Sadness-Neutral* in sACCl and the contrast *delay*_*MW2*_*-delay*_*WM1*_ in the left dlPFC (*R*_*Pearson*_ = – 0.5197, *p* = 0.0132). Moreover, this anticorrelation was significantly stronger (permutation test, *p* = 0.042) in the *high-sadness* group (*R*_*Pearson*_ **=** –0.6639, *p* = 0.0186), than in the *low-sadness* group (*R*_*Pearson*_ = –0.1264, *p* = 0.7278) (Fig. 6B).

We then wondered if the functional connectivity between mFPl and sACCl was also modulated by sadness. Indeed, the connector hub mFPl presented a significant interaction (*high-sadness/low-sadness* vs. *Neutral-WM1/Sadness-WM2*) in its correlation with sACCl (Fig. 6A, right, 3-way ANOVA, *p*_*unc*_ = 0.043). Correlations of mFPl with all other areas did not reach significance (p>0.05). The correlation between sACCl and mFPl showed a marginally significant increase after sadness provocation only in the *high-sadness* group (Fig. 6A, right, paired sample *t*-test *p* = 0.083 for *high-sadness* and *p* = 0.26 for *low-sadness*). The correlation between mFPl and sACCl points at the association of the mFPl-sACCl connection with the sadness-dependent modulation of mFPl network degree (see Fig. 5B).

In summary, we found sadness-related effects at three different levels: behavioral, in functional activity, and in network structure. Importantly, we evaluated the association with sadness by testing the interaction between behavioral paradigm and sadness intensity report, which emphasizes the role of sadness experience in all these modulations. At the behavioral level, the subjects that reported highest emotional scores diminished their performance in the working memory task after sadness provocation (Fig. 2A and Table 3). At the level of functional brain activity, we found an overall decrease in activation in the cognitive areas (Supplementary Fig. S1C), an increase in sACCl activity (Fig. 2B,C) and an inter-individual anticorrelation between sACCl and dlPFCl activity (Fig. 6B). Finally, the graph analysis showed a stronger segregation between emotional and cognitive networks following a strong emotional experience (Table 4), with the connectivity degree of the cognitive connector hub dlPFCl being down-regulated after sadness provocation (Fig. 5A) and that of the emotional connector hub mFPl being up-regulated by sadness intensity (Fig. 5B). Sadness intensity also modulated the functional connectivity of these connector hubs, via sACCl: it increased the correlation between sACCl and mFPl and the anticorrelation between sACCl and dlPFCl (Fig. 6A). We summarize these findings schematically in Figure 7.

**Figure 7:**
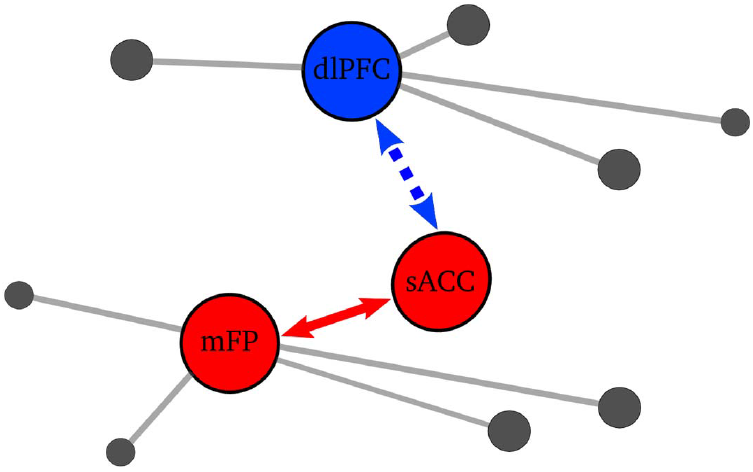
Schematic summary: Sadness-related effects in functional activity and in network structure. The sACC was a key area: following sadness provocation and specifically for high-sadness participants it showed an increase in the negative functional connectivity with dlPFCl and an increase in the positive functional connectivity with mFPl. These functional connectivity modulations could underlie the modulations of dlPFC and mFP as hubs of their respective networks: The dlPFCl degree was decreased after sadness provocation and the mFPl degree was modulated by sadness intensity. Red (blue) denotes *degree increases* (*degree decreases*) and dashed (continuous) arrows denote negative (positive) functional connectivity.

## Discussion

We applied here graph-theoretic network analysis to identify functional changes in brain networks associated with states of sharp emotional and cognitive contrast. Based on functional activations in concatenated working memory and sadness induction tasks, we identified two distributed brain networks, one comprising cognitive regions (dlPFC, IPS, iFG, mSFG, PCG) and another one comprising emotional regions (mFP, sACC, mOFG, Amy, Hip), which could be further decomposed in cortical and subcortical partitions^43^. The modularity of these brain networks increased following sadness induction, consistent with the hypothesis that a sadness state increases the competition between emotional and cognitive subnetworks^17, 44^. In contrast, we found a reduction of modularity within the emotional community^43^, suggesting a role of sadness experience in functionally integrating cortical and subcortical processing within this subnetwork. Two areas emerged from our analysis as task-related connector hubs in the cognitive and emotional brain subnetworks based on their modulation by the sadness induction protocol: the dlPFCl in the cognitive community and the mFPl in the emotional subnetwork. Our data did not support direct interactions between these two hubs but instead coordination via an interposed area, the sACC, as the mediator of sadness-induced modulations in network structure.

In the cognitive network, the connector hub dlPFCl presented a significant decrease in connection degree following sadness induction (Figs. 4C and 5A), suggesting that the sadness state reduced the effective coupling of this area and thus its ability to influence brain processing. Recent evidence has shown that the fronto-parietal brain network, which underlies cognitive control^45, 46^, has especially high global connectivity (i.e., average connectivity with the rest of brain regions^47^), and the global connectivity of the left dlPFC was specifically identified as the mechanism by which the fronto-parietal network might control other networks^48^. Moreover, previous work also attributed a top-down control role to dlPFC in spatial working memory based on neuroimaging data and computational models^49^. Integrating previous literature and our results, the decreases in the degree of the dlPFCl could be related with the overall decrease of the BOLD activity in the cognitive network after sadness experience (Supplementary Fig. S1C) and the decline in the participants’ WM performance (Fig. 2A, Table 3) based on the dlPFC diminished capability of exerting cognitive control during the WM task.

Several studies have found attenuated spatial WM performance during negative task-irrelevant affect^7–10, 12, 50^, associated with marked decreases in dlPFC activity^7, 8, 12, 13^. However, these studies were performed using intervening task-irrelevant aversive stimuli and are thus subject to possible confounds due to attention capture by the noxious stimulus. To address this issue we designed here a paradigm with separate episodes of strong conflicting emotional and cognitive demands (Fig. 1), respectively. In these tasks the outcomes do not depend on the integration of emotional and cognitive aspects. We used a sadness provocation task^29^ to induce a sadness state, followed by a spatial WM task^27^ (*Sadness-WM2* paradigm), and we compared with a control paradigm that concatenates a neutral epoch and spatial WM task (*Neutral-WM1* paradigm) (Fig. 1). In the *Sadness-WM2* paradigm, the cognitive modulations mediated by emotional demands were provoked by an emotional state elicited before the cognitive task. Therefore, unlike previous studies our results do not depend on external distractors or emotional stimuli during the WM task. To our knowledge, only one study before has used a similar strategy on medicated depressed patients^51^. They found WM performance deficits following sadness induction in both controls and depressed patients. This is in line with our finding of impaired WM performance in subjects with higher sadness scores, and supports the role of emotional states in conditioning cognitive function, without any confounds of possible acute attentional shifts by intervening cues as in previous studies.

Intensity and duration are two central characteristics of an emotional response^52^. In our task, participants provided a subjective rating (on a scale 0-7) of the sadness intensity reached after the scanning session. We confirmed (Fig. 2) that this report was indeed measuring effective sadness intensity by validating its correlation with cognitive performance^51^ and with the activation of sACC^29^. Negative emotional traits in healthy people are known also to be associated with increased sACC activation following sadness induction^53^, so our sadness report may be also associated with personality traits of the participants. We then used this report in all our analyses to confirm the unambiguous association of sadness with differences between our two behavioral paradigms, and thus overcome two confounds in our paradigm. For one, in our paradigm we contrasted the induction of an emotional memory with a non-emotional resting epoch without any biographical memory component. We reasoned that this would emphasize the competition between cognitive and emotional processing, while also simulating the rumination associated with depression, but it did incorporate a memory confound that had to be addressed. Secondly, we did not reverse the order of the paradigms, because a previous study found that the sadness block generated some residual effect in control blocks^51^. This could pose interpretation problems associated with the sequence of tasks (practice, tiredness). We addressed both of these confounds in our analyses by testing the relation of our effects with the intensity of the sadness reported by participants. Specifically, we tested the statistical significance of an interaction between the factors *paradigm* and *sadness intensity* in our analyses of variance (ANOVA) tests. Most of the changes in network structure and functional connectivity reported in this study are supported by such a significant ANOVA interaction, thus supporting their unambiguous association with a change in emotional state.

The emotional connector hub was identified as area mFPl based on its modulations by emotional demands. Sadness experience increased the degree of the mFPl in the *high-sadness* group relative to the *low-sadness* group (Fig. 5B), suggesting that intense sadness increases the influence of mFPl on other brain areas. The mFP (part of medial prefrontal cortex) has been described as part of the default mode network, which drives the self-reference processes^54–58^. The modulation in the mFPl degree by sadness intensity could be related with more intense self-reference processes in participants of the *high-sadness* group.

Modulations of the mFPl and dlPFCl degree in our study are in line with the flexible hub theory recently presented^59^ and they suggest that these hubs are capable of functional connectivity adaptations in order to balance cognitive and emotional demands. We found that these adaptations occur coordinated through the sACCl, as it showed more negative functional connectivity with dlPFCl and more positive functional connectivity with mFPl following sadness provocation, and specifically for *high-sadness* participants (Figs. 6 and 7). Remarkably, we also found an inter-individual negative correlation between sACCl and dlPFCl BOLD activity, which was higher in the *high-sadness* group (Fig. 6B). Such result is similar to the inter-individual negative correlation found between the amygdala and inferior frontal gyrus during a working memory task with a negative task-irrelevant stimulus presented during the delay^60^.

There has been substantial debate surrounding the appropriate interpretation of negative correlations observed with resting state functional connectivity when including a preprocessing step termed global signal regression^36–39, 61, 62^. This data processing step can improve the specificity of resting state correlations and the correspondence with anatomy^36^ and electrophysiology^37^, but there are mathematical concerns that anticorrelations could emerge as an analysis artifact^39^. In order to test if global signal regression generated artifactual correlation patterns in our data, we repeated our analyses without global signal regression preprocessing and we found that this did not affect the relative relationships between functional connectivity in our conditions of interest (Supplementary Fig. S4). However, the results with data preprocessed with global signal regression fit together more consistently and provided an easier interpretation. In particular, note that without global signal regression the correlation between sACC and dlPFC became practically zero after sadness induction (Supplementary Fig. S4A), which would be interpreted as sACC and dlPFC becoming decoupled. This decoupling does not fit with other results, in particular with the strong inter-individual correlation that we found between BOLD activity in sACC and dlPFC (see also^18^), especially in the *high-sadness* group (Fig. 6B). Also, a decoupling effect of sadness is at odds with the observed correlation between focal PFC stimulation (TMS) treatment outcome and dlPFC-sACC anticorrelation strength in depressed patients^63^. Because the results are qualitatively unchanged by global signal regression, but provide a much more direct interpretation, we favor here this preprocessing step.

Thus, in our analysis the sACC was not identified as a connector hub area but it did emerge as a key region that coordinates cognitive and emotional connector hub areas, and is thus capable of influencing the global functional network structure. These results provide a new perspective on the previously reported implication of sACC in sadness and depression. Previous studies consistently associate sACC with acute sadness, major depression and antidepressant treatment effects, suggesting a critical role for this region in modulating negative mood states^18, 29, 64^. In addition, sACC connections to the brainstem, hypothalamus, and insula have been implicated in the disturbances of circadian regulation associated with depression and it has been described as a visceral-motor region^65–67^. Reciprocal pathways linking sACC to orbitofrontal, medial prefrontal and various parts of the anterior and posterior cingulate cortices constitute the neuroanatomical substrates by which primary autonomic and homeostatic processes influence various aspects of learning, memory, motivation and reward^65, 68–70^. In depressed patients, the resting-state sACC functional connectivity with the default mode network (DMN) was found stronger than in control participants, and it further correlated with the length of the patients’depressive episodes^71^. All these data reinforce the idea that sACC is implicated in sadness regulation and our results indicate that this could be by means of its regulatory role in relation to two hub network areas, rather than a direct driving mechanism. In our study, participants that reported strong sadness experience had brain activity patterns similar to those previously reported for depressed patients: an activation of the sACC ^17, 33^, a deactivation of dlPFC^33, 72, 73^ and an increase in the anticorrelation between sACC and dlPFC ^18, 63^. This underscores the idea that the network dynamics underlying negative emotional state in healthy subjects could be, when pathologically exacerbated, responsible for behavioral symptoms in depressed patients^17, 18^. Indeed, the three areas that we have identified have been repeatedly associated with antidepressant treatments: response to selective serotonin reuptake inhibitors is related to sACC activity^74–76^, TMS of dlPFC is most effective in sites strongly anticorrelated with sACC^63^, response to TMS of dorso-medial PFC depends on the connectivity of the mFP^77^, response to cognitive behavioral therapy is related to changes in the three areas^76^, and recent results of DBS (subgenual white matter stimulation) link its therapeutic effect to fibers reaching the mFP^20^.

The fact that some of these interventions are very focal and yet address symptoms supported by the activity of system-wide networks^18^, could be explained by our findings of a modular structure in these networks, coordinated by one area (sACC) regulating two connector hubs (dlPFC and mFP). Conversely, also a very focal dysfunction in the network, in one of these coordinating areas, could have profound impact in functions subserved by distant circuits, as recently shown in a computational network model of sACC and dlPFC^44^. Depression as a circuit-level manifestation of a sharply localized disease remains an enticing hypothesis, supported by distributed sadness circuits that interact via very specific coordinating cortical areas.

## Acknowledgments

This work was funded by the Ministry of Economy and Competitiveness of Spain, and the European Regional Development Fund (Refs: BFU2009-09537, BFU2012-34838), by the foundation “La Marató de TV3” (Ref: 091430), and by the Secretaria d’Universitats i Recerca del Departament d’Economia i Coneixement de la Generalitat de Catalunya (Ref: SGR14-1265). JPRM was supported by the Spanish Ministry of Economy and Competitiveness (FPI program). The work was partially carried out at the Esther Koplowitz Centre, Barcelona. We acknowledge MRI assistance from the Dept. of Radiology of Hospital Clínic, Barcelona (Núria Bargalló, César Garrido, and Santiago Soto). We thank Helen Mayberg for a careful reading of the manuscript.

## Author contribution statement

JPRM and AC designed the experiment and the analyses. JPRM and BO recruited participants and collected the data. JPRM and JP performed data analysis. JPRM and AC wrote the manuscript, and PV provided critical review. PV and AC secured funding for the study. All authors approved the manuscript.

## Data availability statement

The datasets generated during and analysed during the current study are available from the corresponding author on reasonable request.

